# Causal event structure shapes convergent reactivation signatures in human cortex and language model

**DOI:** 10.64898/2026.09.28.754904

**Authors:** Zihan Zhang, Yuting Chen, Qiuyi Liu, Xinyi Zhao, Xiao Ding, Xiaoyu Fei, Juntong Du, Yu Bao, Junhao Gao, Xingqi Huang, Bibo Cai, Xia Liang, Bing Qin, Ting Liu

**Affiliations:** Research Center for Social Computing and Interactive Robotics, Harbin Institute of Technology, Harbin, Heilongjiang, China; Shanghai Gener Arfysica Intelligent Healthcare & Brain Science Research Institute, Shanghai, China; Shandong Provincial Key Laboratory of Clinical Research for Pancreatic Diseases, Tumor Immunology and Cytotherapy, Medical Research Center, The Affiliated Hospital of Qingdao University, Qingdao, Shandong, China; Shanghai Innovation Institute, Shanghai, China

**Keywords:** Narrative comprehension, Causal event structure, Neural reactivation, Representational geometry, Persistent homology

## Abstract

Humans adapt to changing environments by understanding the latent causal structure of events, and modern large language models (LLMs) increasingly exhibit analogous abilities. However, elucidating how causal knowledge is represented and integrated in the human brain and in LLMs poses a substantial challenge for cognitive science. To address this, we combine topological analysis of representational manifolds with neuroscientific analyses of reactivation. We find that event-level causal relations modulate temporally structured neural reactivation in the human brain, particularly in the angular gyrus, precuneus and prefrontal cortex. By contrast, LLMs express these relations through positionally structured representational reactivation. Despite these differences, the resulting reactivation patterns exhibit similar intrinsic manifold structure across the two systems. Together, these findings suggest that the human brain and LLM share a common principle for organizing causal knowledge.

## 1 Main

Understanding a narrative requires connecting events beyond their temporal order. An earlier action may establish the conditions for an outcome revealed much later, and understanding that outcome requires relating it to what came before. To support this, the brain parcels continuous experience into discrete chunks of information, termed ‘events’ [1, 2]. These events are not stored independently but are organized into a relational network [3]. Prior work has shown that causal structure is a key organizing principle of this network and that events occupying more central positions within it are more likely to be recalled [4]. However, how earlier events are brought into relation with their outcomes during ongoing comprehension remains unclear.

Event boundaries provide an opportunity for such integration. Neural representations of preceding events are reinstated at boundaries, and stronger reinstatement is associated with better subsequent memory [5, 6]. During naturalistic comprehension, this reactivation extends to remote events sharing the current semantic context [7]; a recent work further reports boundary-related reinstatement of causally related past events [8]. Whether events are causally related, however, does not establish how their roles as causes and outcomes shape the process of linking them. Constructionist accounts propose that readers seek explanations for unfolding events to maintain narrative coherence [9], and behavioural experiments show that earlier causal information can be reinstated when needed to explain a current event [10, 11]. This explanatory demand motivates the hypothesis that **result-event boundaries elicit stronger reactivation than cause-event boundaries**. An outcome can also have multiple antecedents, whose accessibility during comprehension depends on their prior elaboration and availability [12]. This raises a further question: **how do multiple causal antecedents jointly contribute to the representation expressed at an outcome boundary?**

Beyond linking an outcome to its antecedents, narrative comprehension requires representing causal relations that extend across multiple events. An event can be the consequence of earlier events and the cause of later ones, connecting individual cause–outcome relationships into chains with different relational configurations. Evidence that narrative coherence promotes the integration of temporally distant events in hippocampal representations supports considering these relationships at the sequence level [13]. If reinstated antecedents contribute to the representation of an outcome, causal integration may shape both individual event representations and the relationships among them. These relationships define a geometry: each event is represented by a distributed activity pattern, and the relative positions of these patterns describe their similarities and differences. Across a causal chain, this geometry may therefore carry information about how its events are connected. Studies showing that neural population geometry can express abstract task structure, and that complex activity patterns can occupy low-dimensional manifolds, provide a basis for this possibility [14, 15]. We therefore hypothesize that **causal and non-causal event sequences exhibit distinguishable low-dimensional representational configurations, and that different patterns of causal connections are likewise distinguishable in this geometry**.

Comparing human brains with large language models (LLMs) offers a way to test whether this causal organization reflects representational principles shared across biological and artificial systems. Prior work has linked contextual language-model representations to human neural responses [16–18] and identified shared geometric patterns across the two systems [19]. Neural activity unfolds over time, whereas LLM representations are indexed by token positions; both can be examined with respect to the same narrative event sequence (Fig. 1c). At an event boundary, the brain can reinstate activity patterns associated with earlier events, a process we term **neural reactivation** (Fig. 1a,b). In autoregressive transformers, masked self-attention allows information from preceding positions to contribute to later hidden states [20]. The hidden state at the end of an event description can therefore incorporate information from earlier events (Fig. 1d,e). We use **representational reactivation** operationally to describe the measured re-expression of earlier event representations in these later contextual states, as indexed by boundaryto-event representational similarity. We therefore ask **whether causal relations similarly organize the re-expression of earlier event information at outcome boundaries and the geometry of event representations across causal chains in humans and LLMs**.

**Fig. 1.**
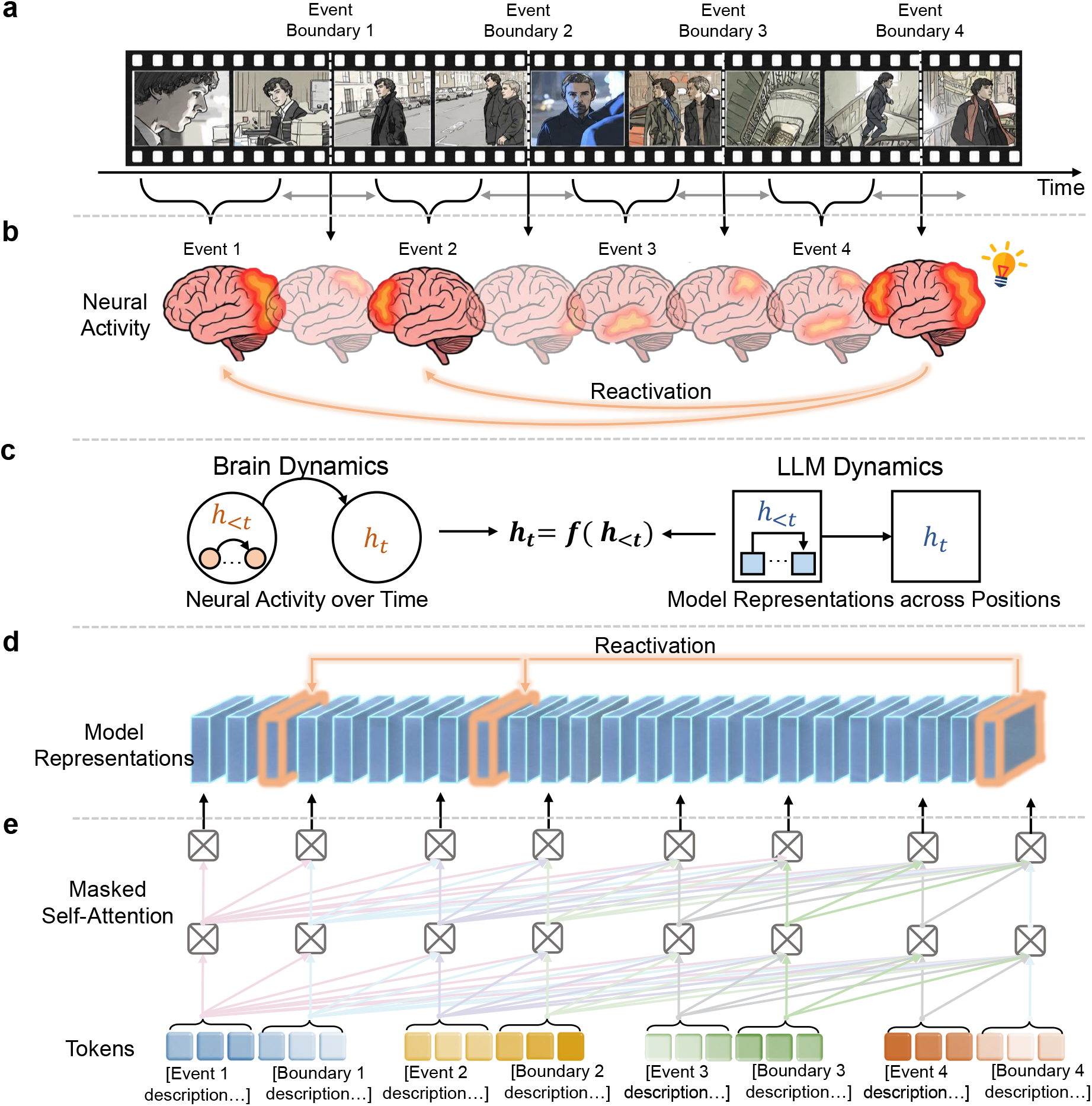
The event-based framework used to compare reactivation dynamics in the human brain and decoder-only LLMs during naturalistic narrative processing. **a**, Participants watched movies during scanning. We defined *event boundaries* as the TRs corresponding to scene transitions. Events were defined as the average activity patterns across all TRs within each scene, excluding TRs located within several time points of the adjacent event boundaries (5 TRs, corresponding to 7.5 s, in Sherlock; 3 TRs, corresponding to 6 s, in Studyforrest) [7]. All analyses accounted for hemodynamic delay. **b**,Reactivation is the reinstatement of past neural activity patterns. **c**,Brain dynamics and LLM dynamics are comparable because both can be expressed within the same sequence-based dynamical framework. **d**,Model Representations across Positions. We use the hidden state at this last position as a compact representation of the corresponding event (or boundary), as it aggregates information from the entire span via self-attention. **e**,Most contemporary LLMs adopt a decoder-only Transformer architecture, in which masking ensures that each token can attend only to preceding tokens.

Here, we combined behavioural measures of narrative retention, neural responses from two independent fMRI datasets, Sherlock [21] and Studyforrest [22], and LLM hidden-state representations elicited by matched event descriptions. We examined causal organization at three complementary levels. At the behavioural level, perceived causal roles were associated with human recall and event mentions in LLM-generated summaries. At the level of representational dynamics, result-event boundaries exhibited stronger reactivation than cause-event boundaries, and event-level reactivation patterns showed reliable cross-system correspondence. To estimate the joint contributions of multiple antecedents, we reconstructed outcome-boundary representations as weighted combinations of preceding event representations and compared the estimated weights with annotated causal strengths. At the sequence level, manifold-derived features distinguished causal from non-causal sequences and differentiated causal-chain configurations. Together, these findings suggest that **causal structure provides a shared organizing principle for sequential representations in human brains and LLMs. It is reflected both in the re-expression of earlier event information at individual boundaries and in the organization of representations across extended sequences**.

## 2 Results

We combined behavioral measures, naturalistic fMRI data and LLM representations to investigate the relationships between causal structure, narrative memory and representational organization. The behavioral experiments in Section 2.1 showed that causally relevant events were preferentially retained in human recall and LLM-generated summaries. To further investigate the causes of this property, in Section 2.2 we analyzed human fMRI data and LLM hidden-state vectors using a boundary-related reactivation approach. We found that reactivation was highly correlated with events causality and showed a high degree of correspondence between the human brain and the LLM. Extending this analysis, in Section 2.3 we used linear reconstruction to characterize the contributions of earlier event representations to boundary representations and found that the resulting event-to-boundary reconstruction matrices aligned with annotated causal relations. Building on these event-level correspondences, in Section 2.4 we compared representational manifolds across complete event trajectories to assess whether the similarities extended to the organization of event sequences. We found that causal and non-causal trajectories differed in their geometry and that humans and the LLM exhibited similar patterns of organization.

### 2.1 Learning behaviour was related to the causal attributes of events

To obtain comparable event-level measures of causal judgement and retention, we implemented parallel protocols for human participants and the LLM (Fig. 2a–c). Participants viewed the Sherlock audiovisual stimulus segmented into 50 events, judged each event online as having no clear bridging function (*Independent*), explaining previous events (*Result*), setting up subsequent developments (*Cause*) or being uncertain (*Other*), and were then asked to summarize the narrative from memory. Using matched text descriptions, the LLM made incremental causal-role judgements according to the same categories and generated a summary after processing the complete narrative. Human recall summaries and LLM summaries were aligned to the same event units, yielding human recall richness and model-side event-mention measures as approximate event-level readouts of narrative retention. Full procedural details are provided in Methods 4.1.

**Fig. 2.**
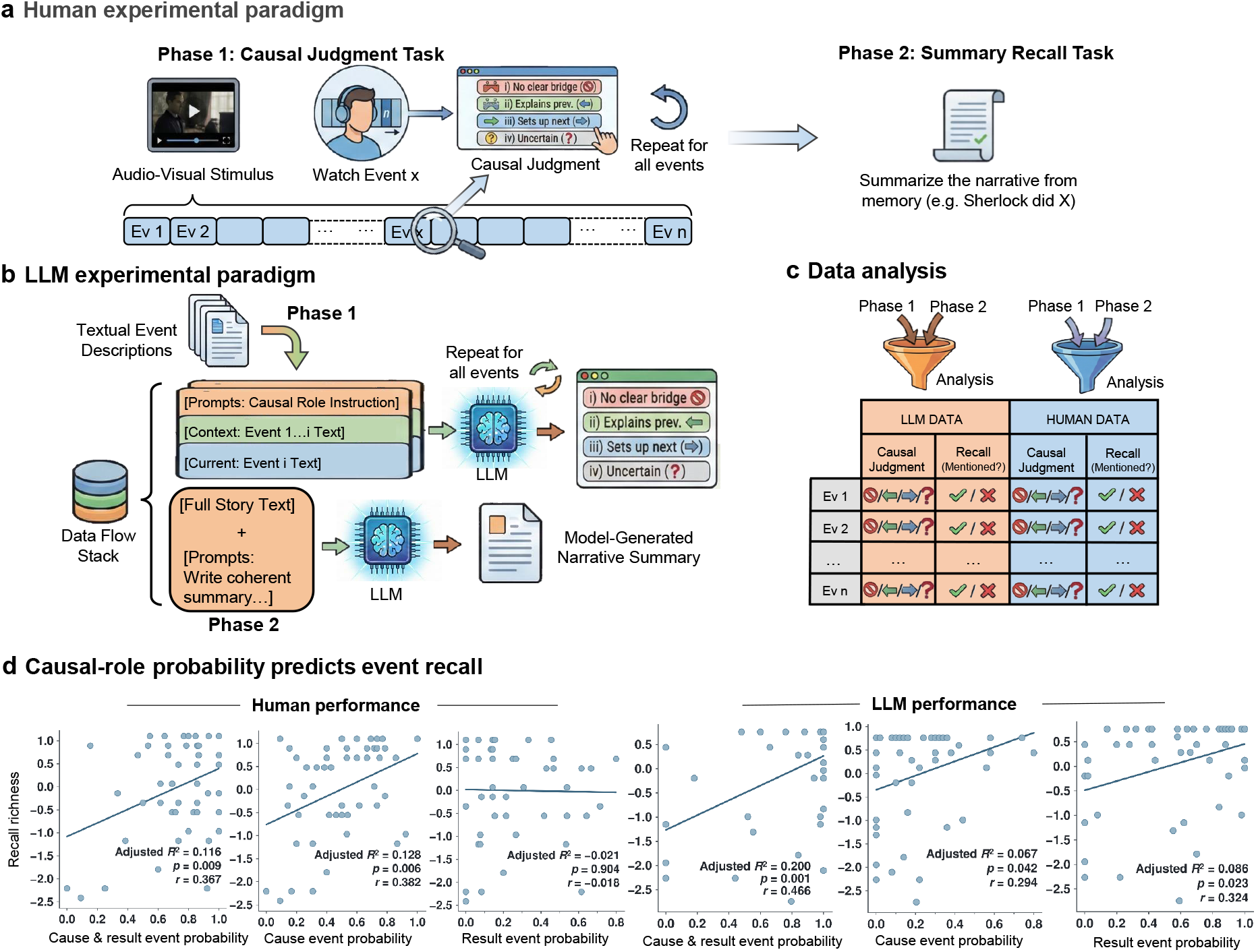
behavioral and LLM paradigms reveal different relationships between causal role and memory. **a**, Human participants viewed a naturalistic movie segmented into predefined event units, made online judgements of each event’s causal role during viewing, and were then asked to summarize the narrative from memory after the movie. **b**, LLMs received matched text-based event descriptions incrementally, generated causal-role judgements for each event using the same response categories, and then produced a global story summary. **c**, Extract causal judgment and recall richness of LLM and humans. **d**, Human event-recall richness as a function of three event-level causal-role measures: combined cause- and-result event probability, cause event probability and result event probability. Each dot denotes one event, and the orange line indicates the fitted linear regression.

Events assigned stronger causal roles were more likely to be recalled by participants and mentioned in LLM-generated summaries. As shown in Fig. 2d, the combined probability that an event was judged as either a cause or a result was positively associated with human recall richness (*r* = 0.367, adjusted *R*^2^ = 0.116, *P* = 0.009) and with its mention in LLM-generated summaries (*r* = 0.466, adjusted *R*^2^ = 0.200, *P* = 0.001). However, the specific causal profiles differed between the two systems. In humans, recall was most strongly associated with the probability that an event served as a cause of subsequent developments (*r* = 0.382, adjusted *R*^2^ = 0.128, *P* = 0.006), whereas result-event probability was unrelated to recall (*r* = ™0.018, adjusted *R*^2^ = ™0.021, *P* = 0.904). This dissociation suggests that human memory preferentially retains events with forward-looking causal potential, rather than events that merely close out prior causal chains. In the LLM, both cause-event probability (*r* = 0.294, adjusted *R*^2^ = 0.067, *P* = 0.042) and result-event probability (*r* = 0.324, adjusted *R*^2^ = 0.086, *P* = 0.023) were positively associated with summary-based retention, indicating a more even weighting of forward- and backward-looking causal roles. Thus, although event causality was positively associated with retention in both systems, humans placed greater emphasis on causally generative events, whereas the LLM’s retention was comparatively balanced across generative and resultant roles.

### 2.2 Causal event structure is associated with boundary-locked reactivation in humans and LLMs

To determine whether event causality was associated with boundary-locked reactivation, we adopted the directional reactivation framework developed by Hahamy et al. [7] for naturalistic-event processing. This framework quantifies reactivation from the temporal asymmetry of boundary-to-event similarity, comparing representations of preceding and following events. The past–future contrast reduces the time-symmetric component of background similarity arising from shared semantic content, character, clue and narrative (Supplementary Information 9). Reinstatement of past events should preferentially increase past-directed similarity. Following the original framework, we excluded temporally adjacent matrix entries to reduce local temporal autocorrelation and used symmetric non-boundary control time points in the whole-brain localization analyses to assess boundary specificity (Methods, Section 4.3; Supplementary Information 3).

We then tested whether this directional reactivation signal varied with event-level causal role. Reactivation indices were compared across four behavioral categories: *Result, Cause, Independent* and *Other* (Fig. 3a,b).

**Fig. 3.**
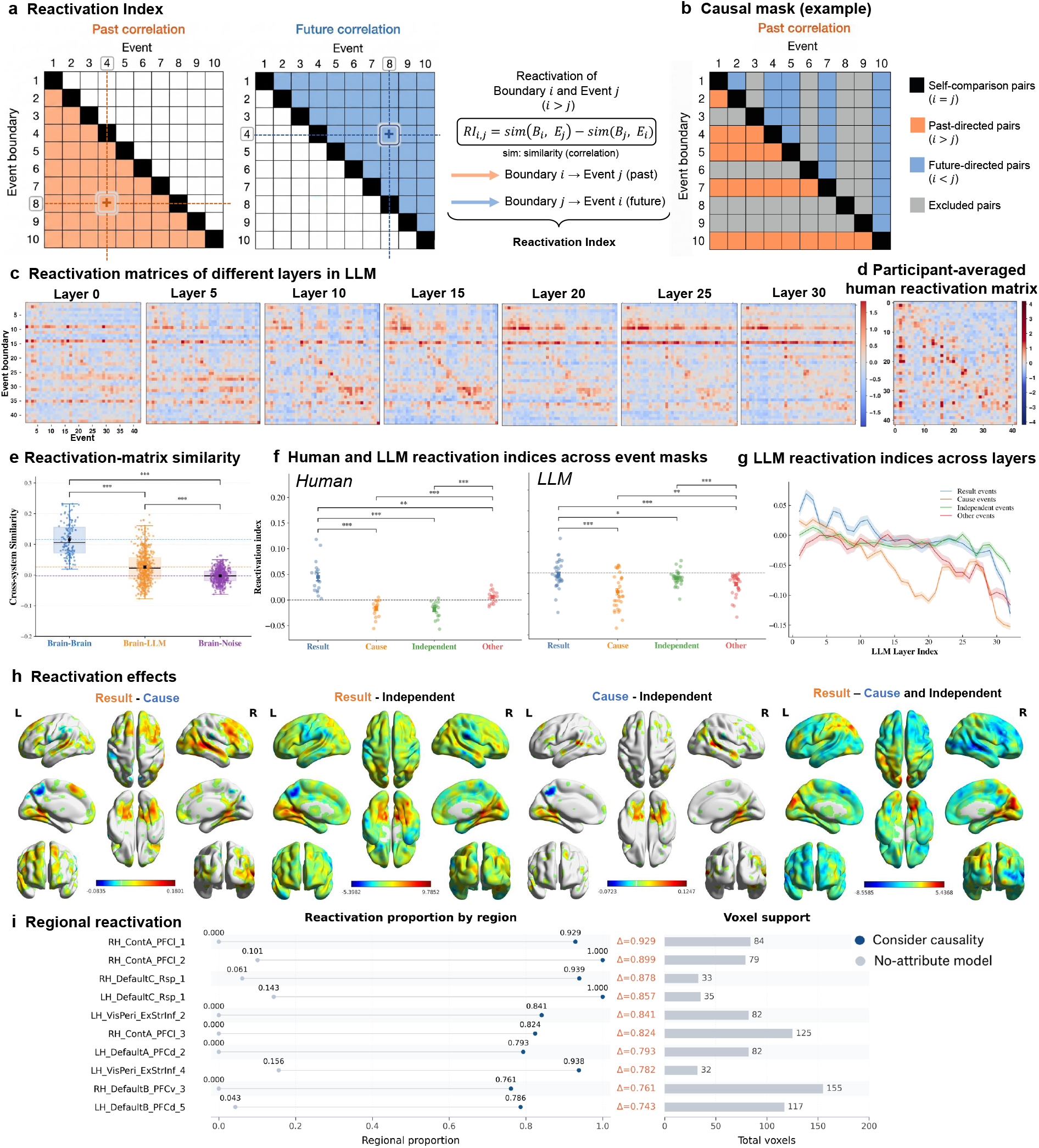
Event-boundary reactivation tracks causal structure in the human brain. **a**, Schematic illustration of the reactivation analysis. **b**, Mask used to test the effect of event-level causal structure on the reactivation matrix. **c**, Reactivation-strength matrices (standardized) for representative layers of the LLM. **d**, Group-averaged reactivation-strength matrix (standardized) in the human brain. **e**, Brain–LLM similarity analysis revealed reliable correspondence between human and model reactivation structure beyond permutation-based null controls. **f**, Reactivation indices computed under causal-role masks in humans and across LLM layers. Both systems showed maximal reactivation at Result events, but the effect was substantially stronger in humans. **g**, Reactivation indices across LLM layers. **h**, Human cortical reactivation-index contrasts between event causal roles. dRI denotes the across-participant mean of within-participant RI differences. The final map shows the intersection of same-direction Result–Cause and Result–Independent effects. Only significant voxels are displayed ((n=17); two-sided paired (t)-tests, (*p <* 0.05); Result–Cause FWE-corrected, other maps uncorrected). **i**, Regional comparison of the Result–Cause map (blue) with the original reactivation analysis (grey). Shown are the ten Schaefer 400-atlas parcels with the largest positive differences in significant-voxel proportions. Dots indicate proportions, orange Δ labels indicate their differences (blue minus grey), and bars show total voxel counts per parcel.

#### Reactivation structure in the human brain and LLMs

We first examined the overall reactivation structure independently of causal category. The reactivation-strength matrices of the LLM and the human brain exhibited a corresponding triangular asymmetry (Fig. 3c,d). This triangular asymmetry is a characteristic signature of retrospective reactivation, reflecting the preferential reactivation of preceding events by subsequently encountered events. The complete reactivation-strength matrices for all LLM layers and human participants are provided in Supplementary Information 4. In humans, mean coupling was higher in the *Past Correlation* triangle than in the *Future Correlation* triangle (mean coupling 0.043 versus ™0.058; paired comparison across 17 participants: *t*(16) = 5.6, two-sided *P* = 4 ×10^*™*5^). Thus, boundary–event similarity was directionally structured rather than symmetrically distributed across the matrices. In the LLM, the magnitude of this asymmetry varied across layers and was most pronounced in the earliest layers (Fig. 3c). Later-layer matrices descriptively exhibited more prominent banded structures, characterized by high similarity to a restricted set of event representations. One possible explanation is a gradual transition towards representations increasingly shaped by the model’s evolving output predictions [23–26], though the present data are descriptive and do not directly test this mechanism.

#### Correspondence between human and LLM reactivation matrices

Given the similar organization of the matrices in Fig. 3c,d, we next quantified their cross-system correspondence (Fig. 3e). Mean matrix similarity was 0.117 for Brain–Brain comparisons, 0.025 for Brain–LLM comparisons and ™0.004 for Brain–Noise comparisons. Brain–LLM similarity was higher than Brain– Noise similarity (*P <* 10^*™*3^), indicating reliable cross-system correspondence beyond the noise baseline. Brain–Brain similarity was higher than both Brain–LLM similarity (*P <* 10^*™*3^) and Brain–Noise similarity (*P <* 10^*™*3^). Thus, reactivation matrices shared systematic event-level structure across humans and the LLM, although within-brain correspondence remained substantially stronger than cross-system correspondence.

### Causal-role-specific reactivation in humans and LLMs

Having established that reactivation matrices carry shared structure across brain and LLM, we next examined whether this structure reflects a common organizing principle rooted in the causal role of each event boundary.

Reactivation indices differed across causal-role categories in the human brain, as assessed using a oneway repeated-measures ANOVA (Fig. 3f; *F* (3, 48) = 35.59, *p* = 2.95× 10^*™*12^). Reactivation was strongest at Result-event boundaries and was significantly greater than at Cause-event boundaries (*t*(16) = 7.20, Bonferroni-corrected *p* = 1.27 ×10^*™*5^), independent-event boundaries (*t*(16) = 7.01, Bonferroni-corrected *p* = 1.77 ×10^*™*5^) and Other-event boundaries (*t*(16) = 4.54, Bonferroni-corrected *p* = 2.01 ×10^*™*3^), consistent with a stronger retrospective bias when a causal outcome occurred.

Whereas the human analyses used participants as independent units of inference, the 32 layers of a single LLM constituted dependent measurements within a fixed model. We therefore assessed causal-role differences in the LLM using an event-level restricted permutation test, with an omnibus statistic quantifying differences among the four layer-averaged category means (Fig. 3f,g). Each permitted event-label permutation was applied identically across all layers, and the category-specific masks were recomputed at each iteration, preserving the correspondence of event representations across layers. This analysis revealed an overall causal-category effect (*p*_perm_ = 0.004; 10^3^ permutations). As in humans, reactivation indices were greater at Result-event boundaries than at Cause-event boundaries, with a mean within-layer Result–Cause difference of 0.046 (Bonferroni-corrected across the 6 pairwise category comparisons).

Besides, the standardized Result–Cause difference was greater in the human brain than in the LLM (1.90 versus 1.13; *p* = 9.3× 10^*™*3^). Thus, causal outcomes were associated with retrospective integration in both systems, but this association was more pronounced in the human brain. This difference may help explain the behavioral dissociation in Fig. 2d, whereby human recall was preferentially associated with causal antecedents, whereas LLM summaries weighted cause and result events more evenly.

#### Spatial distribution of causal reactivation in the human brain

To investigate how event causal roles were associated with the spatial distribution of reactivation, we applied the mask illustrated in Fig. 3b to the reactivation matrix. The primary Result–Cause contrast tested whether Result-event boundaries were associated with stronger past-directed reactivation than Cause-event boundaries. For each pairwise contrast, dRI denotes the across-participant mean of the within-participant difference in reactivation index (*n* = 17). Significant Result–Cause differences were observed in the precuneus, prefrontal cortex and angular gyrus (Fig. 3h; two-sided paired *t*-tests, *p <* 0.05, family-wise error FWE-corrected). Positive dRI values in this contrast indicated stronger past-directed reactivation at Result-event boundaries.

Additional Result–Independent and Cause–Independent comparisons were evaluated using two-sided paired *t*-tests at uncorrected *p <* 0.05. The final map in Fig. 3h shows regions where Result events differed significantly from both Cause and Independent events in the same direction, with each comparison meeting the uncorrected *p <* 0.05 threshold. Algorithmic details are provided in Supplementary Information 3.

To characterize the spatial differences associated with introducing event causal roles, we compared the Result–Cause map with our replication of the original reactivation analysis of Hahamy et al. [7], which did not distinguish events by causal role. Within each parcel of the Schaefer 400-parcel atlas [27], we calculated the proportion of significant voxels in each thresholded map and the difference between these proportions (Fig. 3i). The ten parcels with the largest positive differences were located primarily in the retrosplenial cortex, dorsal and lateral prefrontal cortex, and peripheral visual cortex. This comparison descriptively summarized differences in regional coverage between the two maps; no additional inferential test between the analyses was performed.

An exploratory NeuroSynth comparison showed a modest spatial correlation between the Result– Cause map and the meta-analytic map associated with “navigation” (*r* = 0.107) [28], providing descriptive functional context for this spatial pattern.

Together with the preceding analyses, these findings show that event causal roles were associated with the organization of reactivation in both the human brain and the LLM. The two systems exhibited matrixlevel correspondence and enhanced reactivation at causal outcomes, while differing in the magnitude and regional or layer-wise expression of these effects.

### 2.3 Reconstruction weights track causal dependencies across systems

Having established that reactivation was strongest at Result-event boundaries, we next asked whether it reflected the reinstatement of specific causal antecedents. We therefore extended the category-level analysis to individual event pairs: each boundary representation was jointly reconstructed from preceding event representations, and the resulting weights were compared with the annotated causal strengths linking those preceding events to the completed event (Fig. 4a).

**Fig. 4.**
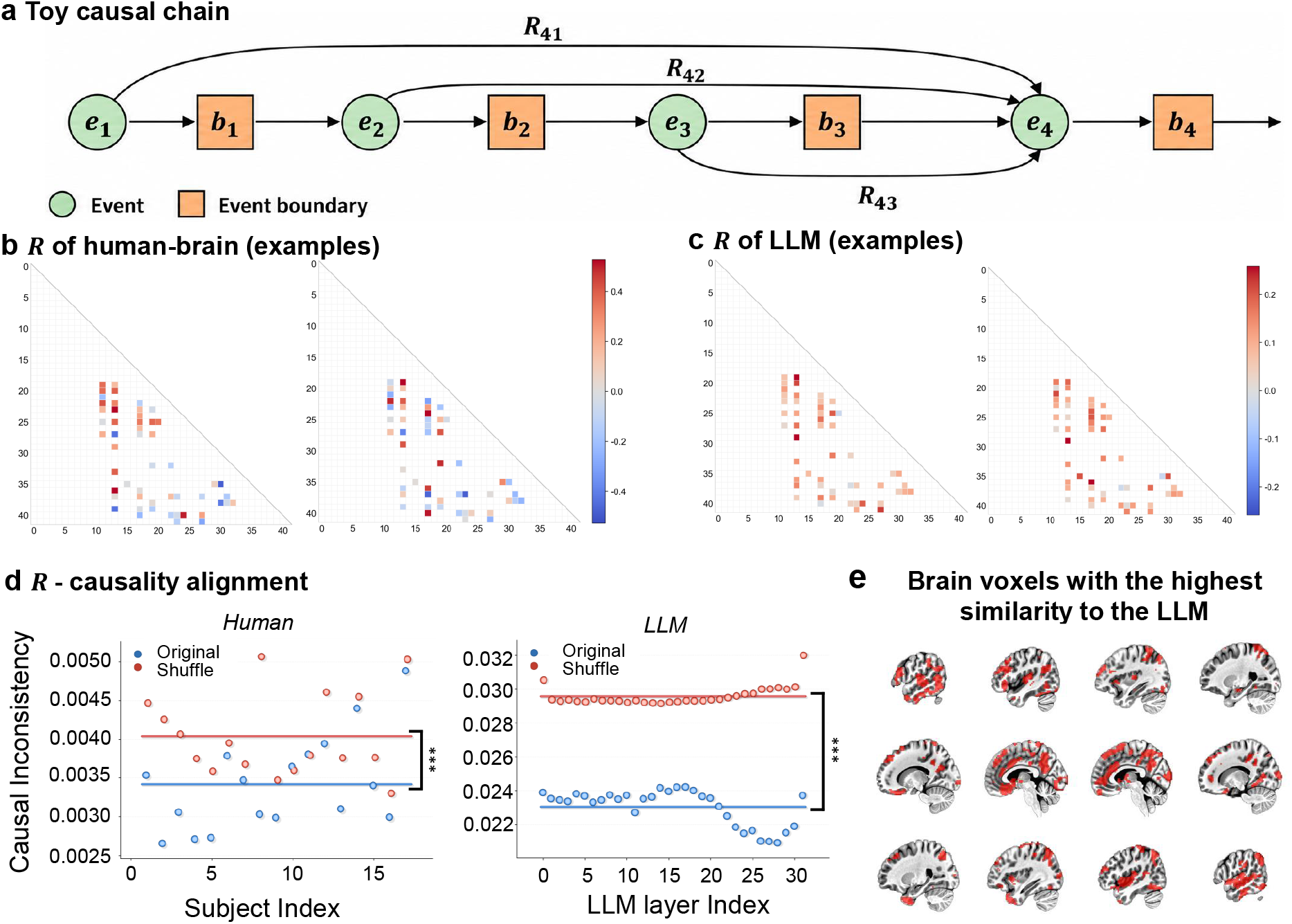
**a**, Toy causal chain used to define event-level causal dependencies. **b**, Example event-to-boundary reconstruction matrixs from human fMRI data. **c**, Example event-to-boundary reconstruction matrixs from LLM hidden states. **d**, Event-to-boundary reconstruction matrices in humans and LLMs exhibited significantly stronger alignment with the weighted causal annotations than expected under random permutations. Dots denote individual participants or LLM layers; horizontal lines denote group means. Statistical significance was assessed against the shuffle baseline. **e**, Cortical voxels showing the strongest similarity between human event-to-boundary reconstruction matrixs and LLM event-to-boundary reconstruction matrixs.

For example, the boundary representation of event 4 is modelled as

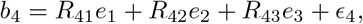

where *e*_1_, *e*_2_, *e*_3_ are the event representations, *b*_4_ is the boundary representation, and *ϵ*_4_ is the residual. The coefficients were estimated jointly using ordinary least squares, minimizing the squared error in reconstructing each boundary representation from preceding event representations. Throughout the reconstruction analysis, *R*_*ij*_ denotes the coefficient relating event *j* to boundary *i* (Methods, Section 4.5). We applied the same model to individual human participants and LLM layers, yielding the reconstruction matrices shown in Fig. 4b,c. In the displayed examples, the human matrices showed larger positive coefficients and negative coefficients closer to zero than the LLM matrices.

#### Correspondence between reconstruction coefficients and causal annotations

We next examined whether these reconstruction weights corresponded to the annotated strengths of causal relations. We computed the distance between each event-to-boundary reconstruction matrix, **R**, and the corresponding weighted causal reference matrix, **C**_weight_, which encoded the graded extent to which one event was judged to influence another. Here, **R** was estimated from fMRI and LLM representations, whereas **C**_weight_ was derived from annotations of the strength of causal relations between events, providing an reference for their objective causal structure. This analysis therefore tested whether the estimated contributions of preceding event representations aligned with the causal structure of the narrative. The weighted causal reference matrices used in this analysis, together with the underlying event-level causal annotations, are provided in Supplementary Information 5.

In both humans and LLM, reconstruction matrices showed smaller distances to the weighted causal reference matrix than the mean distances obtained from 1,000 shuffled reference matrices (Fig. 4d, *P <* 10^*™*3^). Shuffled references were generated by randomly permuting matrix entries within the strictly lower triangle, preserving temporal directionality and the distribution of causal weights. These findings provide evidence that reactivation weights were more strongly aligned with event-level causal structure than expected under random permutations. These findings provide evidence that boundary-related representational reinstatement is organized around event-level causal structure in both systems.

#### LLM-like reactivation patterns map onto human association cortex

Finally, we examined which cortical voxels contributed most to the correspondence between human and LLM event-to-boundary reconstruction matrices. We removed each voxel separately, recomputed the human reconstruction matrix, and measured the change in its discrepancy from the LLM matrix. Voxels with the largest positive changes were selected, up to a 20% voxel budget (Fig. 4e). Their removal produced the greatest loss of brain–LLM alignment. The selected voxels qualitatively resembled the cortical pattern in Fig. 3h, linking the localization of alignment contributions to the preceding reactivation analysis. Together, these analyses provide converging evidence that both human brains and LLMs organize reactivation according to the causal structure of unfolding events, while also enabling a direct comparison of causal-event representations across biological and artificial systems.

Together, the category-level effect in Section 2.2 and the event-pair-level alignment here suggest that reactivation is enhanced at Result-event boundaries and preferentially reflects the specific causal antecedents of the completed event.

### 2.4 Manifold similarity underlies similar reactivation patterns

To further examine how event representations evolve as causal structure unfolds, we defined a causal-chain configuration as the pattern of causal links among an ordered sequence of events. The three- and four-event designations refer to the number of constituent events defining the causal-chain configuration, rather than the number of points in its representational trajectory. For each configuration instance, events were selected from the narrative interval between its first and last constituent events, in chronological order. Non-constituent events with annotated causal links to any constituent event were excluded. The number of trajectory points therefore depended on the narrative interval and event selection, and was not fixed at three or four. The representation of each successive event constitutes a point in the original highdimensional representational space of the brain or LLM. Connecting these points into a trajectory whose successive states and transitions describes temporal evolution in the brain and positional evolution in the LLM. We therefore characterized the intrinsic geometry and topology of these causal-chain trajectories directly in their representational spaces and compared the resulting manifold statistics across the two systems. Here, we also controlled for various possible confounding factors (Supplementary Information 9).

For this analysis, each event pair was provisionally classified as causal or non-causal by several high-performing proprietary LLMs. These annotation models were distinct from the open-source LLM analysed in the main experiments, and disagreements were adjudicated by three human raters through majority vote. The resulting pairwise labels were then combined to identify distinct causal-chain configurations. This procedure yielded three configurations for three-event sequences and eight for four-event sequences (Fig. 5a). Full details of the annotation models, prompts, adjudication procedure and agreement statistics are provided in Supplementary Information 5.

**Fig. 5.**
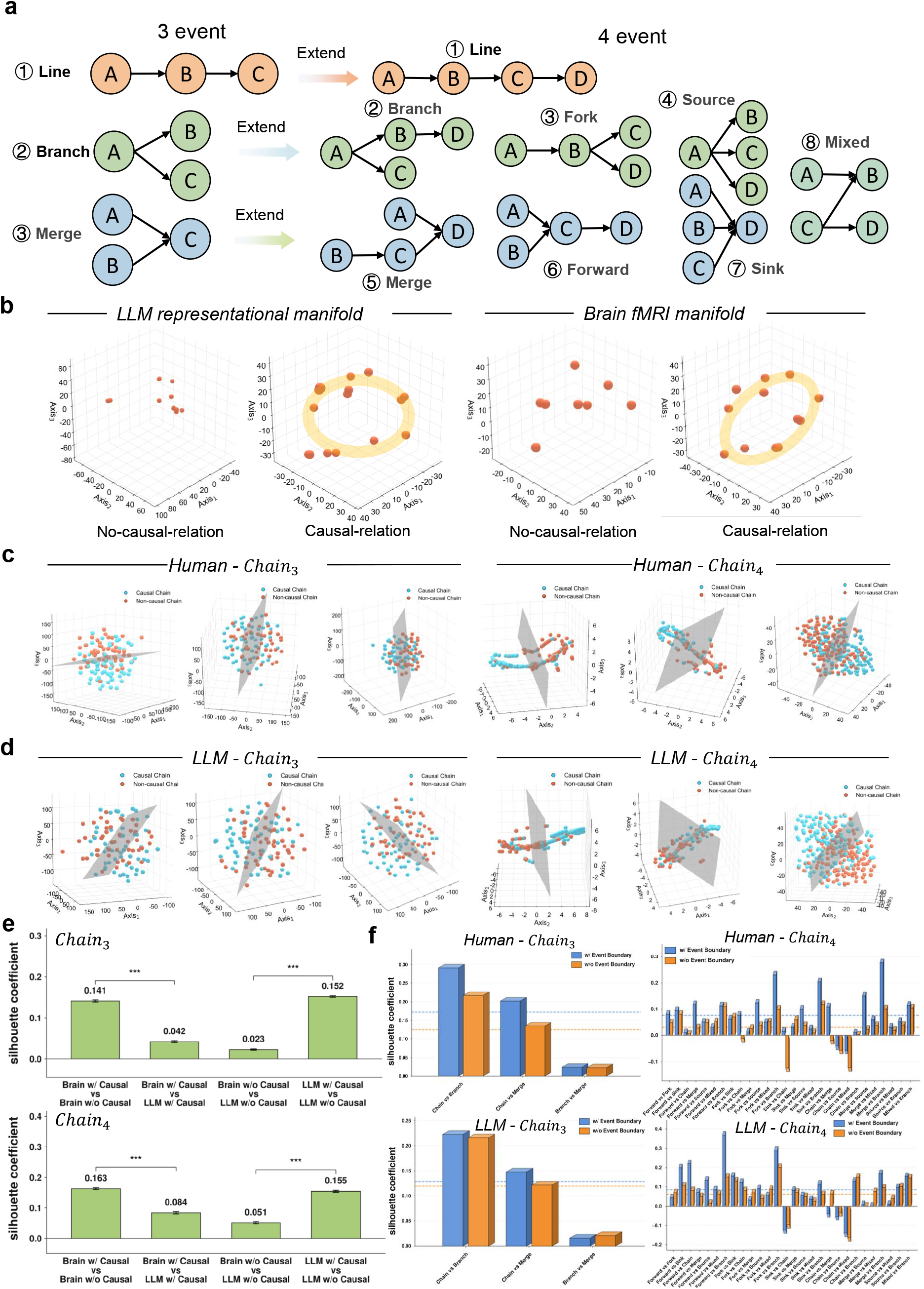
**a**, Annotated causal-chain motifs: three types for three-event chains and eight for four-event chains. **b**, Low-dimensional visualizations of LLM and human representational trajectories (single subject or single layer). PCA/UMAP projections are shown for visualization only; all manifold statistics and persistent-homology analyses were computed from the original high-dimensional representations before dimensionality reduction. **c**,**d**, Low-dimensional visualizations of the topological feature vectors derived from persistent homology. Each point represents one causal chain. **e**, Cross-system comparison of manifold statistics across causal and non-causal conditions. The silhouette coefficient is used here as a measure of system separability. Values closer to zero indicate weaker system-wise separation. **f**, Discriminability of manifolds across distinct causal chains. Blue and orange indicate boundary-retained and boundary-removed comparisons, respectively, with lines denoting the corresponding means. LLM manifold statistics were derived from layers 6–11 for three-event chains and layers 17–22 for four-event chains.

#### Intrinsic manifold analysis and low-dimensional visualization

To characterize the representational geometry of causal-chain processing, we embedded fMRI activity patterns from the human brain and hidden-state vectors from the LLM in their respective high-dimensional state spaces. Then representational geometry was analyzed by persistent-homology method. For visualization, the response vectors were reduced to six principal components and subsequently mapped into three dimensions using UMAP (Fig. 5b). Three- and four-event causal configurations were identified from the pairwise annotations described above. To construct non-causal controls, we randomly sampled starting positions and extracted event sequences of the same length as the causal chains. Candidate sequences containing an annotated causal chain were excluded.

Further, to characterize the topology of representational trajectories, we summarized the standardized high-dimensional responses using persistent-homology features derived from *H*_0_ and *H*_1_, yielding a 450-dimensional topological representation for each trajectory. PCA visualization revealed an apparent separation between causal and non-causal sequences in representative human participants at both chain lengths (Fig. 5c), indicating differences in their fMRI-derived topological organization.

A similar separation was observed in the LLM (Fig. 5d). For each chain length, we identified the three consecutive layers with the highest mean silhouette coefficient computed in the original topological feature space, corresponding to layers 7–9 for three-event chains and layers 20–22 for four-event chains. This shift toward deeper layers for longer causal chains suggests that more complex causal structures are preferentially represented at later stages of model processing. Results across all participants and LLM layers are provided in Supplementary Information 6.

These results provide a geometric account of why both systems exhibit reactivation-like signatures during causal understanding. In the absence of causal relations, fMRI activity patterns and LLM hidden-state vectors showed scattered distributions, suggesting that event representations were weakly organized and did not follow a stable trajectory. By contrast, causal event relations produced a pronounced geometrically regular organization in both systems. These findings suggest that causal structure constrains the geometric organization of neural and model representations in their respective high-dimensional spaces. This regularities reflect structured covariation across successive representational states, with causally linked events aligned along a shared trajectory.

Therefore, the alignment on the manifold shape provides a mathematical basis for reactivation: when a later event boundary falls near to earlier causally linked events on the same manifold, it can reinstate components of those earlier representations, thereby producing the boundary-to-event similarity structure observed in the reactivation-strength matrices.

#### Manifold discrimination under different conditions

We next examined how manifold statistics varied with causal status and system identity, separately for three- and four-event sequences (Fig. 5e). Within each system, causal and non-causal sequences showed separation in their manifold statistics: silhouette coefficients were 0.141 and 0.163 in humans and 0.152 and 0.155 in the LLM, respectively. By comparison, cross-system separation was weaker when causal status was matched. Human–LLM comparisons yielded coefficients of 0.042 and 0.084 for causal sequences and 0.023 and 0.051 for non-causal sequences (Silhouette Coefficients closer to zero indicating stronger consistency). Cross-system comparisons with mismatched causal status also yielded higher coefficients than the corresponding matched comparisons.

This indicate that the extracted manifold statistics retained differences between causal and noncausal sequences within each system, while showing relatively weak separation between systems under the same causal condition. Thus, the observed cross-system correspondence coexisted with sensitivity to causal status. Together, these results support a shared condition-dependent organization of representational geometry in humans and the LLM, with causal event structure constituting a relevant organizing dimension in both systems.

Building on the causal versus non-causal discrimination, we further examined whether manifold statistics could distinguish different causal-chain patterns within the causal condition. Specifically, we compared the manifold statistics associated with the distinct causal-chain motifs shown in Fig. 5a. The results are presented in Fig. 5f.

Specifically, for three-event causal chains, the results revealed a striking correspondence between the manifolds derived from human brain activity and those derived from the LLM. Notably, even after removing the boundary-based contrast, both systems retained an organized manifold structure across events, although the boundary-related features of the manifolds became less pronounced. This suggests that reactivation is not confined to event boundaries, but is distributed throughout event processing; however, event boundaries support a more complete expression of causal structure. The reduction in manifold discriminability after boundary removal was more pronounced in humans than in the LLM, indicating that human event boundaries carry stronger reactivation signals. This observation is consistent with the conclusion drawn from Fig. 3f. Moreover, the mean causal-discrimination effect in the human brain was markedly stronger than that in the LLM, suggesting that the brain more robustly separates relatively simple causal structures.

For causal chains of length 4, the results further demonstrate a strong correspondence between manifolds in the human brain and the LLM. Unlike the case of length-3 causal chains, however, the mean causal discrimination effect in the human within-event manifold after removal of the event-boundary component was closer to zero, suggesting that the representation of more complex causal chains in the brain depends predominantly on boundary-related reactivation. Moreover, causal discrimination in the human brain, both in the presence and absence of the boundary component, was lower than that observed in the LLM. This finding indicates that the LLM is more sensitive than the human brain to relatively complex causal relationships, in direct contrast to the pattern observed for causal chains of length 3.

### 2.5 Replication in the independent Studyforrest dataset

To test whether the findings extended to a different narrative and participant cohort, we conducted parallel analyses using human fMRI data from Studyforrest and LLM representations of the corresponding *Forrest Gump* narrative. These analyses reproduced the principal patterns observed in Sherlock.

Both systems showed preferential reactivation at Result-event boundaries, with human effects involving association cortex and human–LLM reactivation-matrix similarity exceeding the brain–noise baseline (Supplementary Figs. S11 and S12). Event-to-boundary reconstruction matrices also showed closer correspondence to the weighted causal reference than to the shuffled baseline in both systems (Supplementary Fig. S14).

For three-event chains, causal and non-causal sequences again differed in their representational geometry in both humans and the LLM. We did not analyse four-event chains because too few eligible instances were available to support reliable representational analyses. Discrimination between these sequences generally decreased when boundary-related representations were removed (Supplementary Fig. S13). Together, these parallel results extend the association between event-level causal structure and boundary-related representational organization to an independent narrative and participant cohort. Detailed methods, results and figures are provided in Supplementary Information 8.

## 3 Discussion

Understanding causality is central to adaptive behaviour: intelligent systems must encode events and infer how they explain and predict later developments. Here we show that event-level causal structure is reflected in representational dynamics in the human brain and LLMs. In humans, causal antecedents were reinstated at event boundaries in association cortex, including the angular gyrus, precuneus and prefrontal regions. In LLMs, hidden-state representations showed reactivation-like boundary-to-event similarity and causal-chain-dependent representational organization. These findings suggest that causal structure provides a common organizing principle for sequential representations across biological and artificial systems.

In our behavioral experiment, stronger causal roles were associated with greater event inclusion in human recall and LLM summaries. This agrees with classical findings that events in causal chains, especially those with more causal connections, are better remembered, judged more important and more often summarized [29, 30]. Recent work also shows that causal coherence improves memory for the order and relevant details of unfamiliar visual events [4]. Our results extend these findings by distinguishing the contributions of cause and result events. Human recall was associated with events that set up subsequent developments, whereas LLM summaries reflected both causes and outcomes. Thus, causal relevance contributes to memory in both systems, while the particular role an event plays helps determine which information is retained.

Our neural results extend previous work on event segmentation and memory reactivation by estimating the contributions of earlier event representations to activity patterns at event boundaries and showing that these reconstruction weights align with annotated causal structure. Earlier studies demonstrated that event boundaries trigger reactivation of preceding events, supporting the integration of temporally separated events [5, 7]. More recent intracranial recordings also revealed increased hippocampal ripple-like activity at movie boundaries [31]. In our study, human association cortex preferentially reinstated antecedent representations when a causal outcome occurred. The involvement of the precuneus, angular gyrus and prefrontal regions also accords with evidence linking these association regions to narrative integration [32], causal judgement and asymmetric cause–effect representations [33, 34], and episodic retrieval [35, 36]. In a recent study using a narrative with interleaved storylines, causal relations between events were found to predict recall organization and boundary-related reinstatement in the default mode network [8]. Our study further distinguishes the roles of cause and result events and compares reactivation patterns and causal-chain geometry between humans and LLMs. These findings support the view that event boundaries serve as moments of model updating [2], and further suggest that causal relations help determine which earlier information enters the updated model. Together with the behavioral findings, this suggests that cause events provide information worth retaining, while result events provide cues for bringing that information back into current processing.

The LLM comparison offers a computational perspective on causal understanding. In humans, transient input creates pressure to process information rapidly and selectively maintain or retrieve it as new input arrives [37]. Reactivation supports this reuse, consistent with evidence that human replay can reorganize experience according to learned relationships [38]. LLM self-attention instead gives the current position access to earlier token representations within the available context [20]. Attention can be understood as an associative-memory update [39], and induction-head circuits illustrate how transformers locate earlier tokens and reuse the information that follows them [40]. These mechanisms plausibly explain the reactivation-like patterns in LLM hidden states. Both systems can thus connect causally related events through different ways of maintaining and accessing prior information. These differences may help explain the emphasis on antecedents in human recall and on both causes and outcomes in LLM summaries.

In this sense, AI learning may be viewed as a form of data selection, and the natural environment itself can be regarded as an immense structured dataset from which biological intelligence has been selected. From the perspective of convergent evolution, similar computational pressures may lead biological and artificial systems to develop analogous, though not identical, representational strategies.

Besides, a key contribution is to examine this correspondence through representational geometry. Building on studies showing that language-model features predict neural responses [16, 18], we compared the collective organization of causal-chain representations. This connects to evidence for map-like representations of abstract relationships in the hippocampal–entorhinal system [41] and computational accounts of how relational structure is reused across experiences [42]. A causal chain links events within a common explanation, through successive dependencies or multiple causes contributing to one outcome. Comparing chain geometry therefore examines how these related events are represented together. Apparent separation between causal and non-causal sequences in both systems suggests that causal relations are reflected in their collective representational organization. Such organization could help familiar causal patterns guide understanding of narratives with new characters and settings. This manifold-based view links neural dynamics and model representations to the generalization of causal knowledge.

Limitation should be noted. Our analyses focus primarily on reactivation and representational geometry during ongoing narrative comprehension. How these patterns are maintained or reorganized over longer retention intervals remains to be examined. Future work could investigate how repeated experience and memory consolidation shape the persistence of causal event representations.

This study identifies causal structure as a shared organizing principle in humans and LLMs. Neural reactivation and model attention provide different routes for bringing past information into current processing. Similarities in reactivation and causal-chain organization suggest common demands from integrating causes and outcomes. Linking causality, memory, reactivation and geometry provides a framework for understanding how intelligent systems organize experience into coherent accounts of the world.

## 4 Methods

### 4.1 Behavioral Experiment

#### Participants

Fifteen healthy adults participated in the behavioral study (7 men, 8 women; age range 20–29 years). The behavioral cohort was similar in size to the original fMRI cohort. All participants provided written informed consent before participation. The study was approved by the Harbin Institute of Technology Medical Ethics Review Committee (HIT-2025064). All recruited participants completed the full experiment and received monetary compensation (50 RMB per participant). The behavioral dataset will be made publicly available upon publication of the associated work.

##### Stimulus and event segmentation

We used an edited naturalistic narrative stimulus matched to the structure of the *Sherlock* viewing–recall paradigm. The movie was segmented into 50 consecutive events of variable duration (from the original dataset), defined *a priori* and fixed across participants. Each event boundary was followed by a 5-s blank screen that served as a response window for online behavioral annotations.

#### Procedure

The experiment comprised two phases. In Phase 1, participants watched the full stimulus once. Immediately after each event, during the 5-s blank interval, they answered a forced-choice question assessing the causal role of the just-viewed event in the unfolding plot:

Which option best describes the core function of the event you just watched in the story?

Options include: (i) the event does not yet show a clear bridging function (neither a clear cause nor a clear consequence); (ii) the event primarily explains a previous event (consequence/explanation); (iii) the event does not explain earlier events but clearly sets up later developments (cause/foreshadowing); and (iv) uncertain. Participants were instructed to respond before the end of the 5-s window; missed responses were coded as missing. In Phase 2, immediately after movie viewing, participants completed a written summary recall task. They were instructed to summarize the narrative from memory without prompts and were encouraged, but not required, to follow the story’s chronological progression.

#### behavioral measures

For each event, we computed the proportion of participants selecting each causal-role category. The *causal-role probability* of an event was defined as the number of participants who classified that event as either a cause-like or consequence-like event divided by the total number of participants. Recall transcripts were segmented into short clauses or utterances and aligned to the predefined 50 events by two independent coders blind to the causal-role ratings. Disagreements were resolved by discussion or adjudication by a third coder. From this alignment, we derived an event-wise recall measure, termed *recall richness*, defined as the number of participants who recalled a given event divided by the total number of participants.

#### Dawid–Skene aggregation of causal-role labels

To obtain robust event-level causal annotations from multiple noisy individual judgements, we used the Dawid–Skene model for categorical label aggregation. In this model, each participant provides an observed label for each event, while the latent “true” label is unobserved. The model estimates (i) an event-wise posterior probability over categories and (ii) a participant-specific confusion matrix capturing systematic response tendencies, using expectation–maximization. We applied Dawid–Skene to the 15 participants’ event-wise labels to compute a posterior distribution *p*(*y*_*e*_ = *c*) for each event *e* and category *c*.

### 4.2 LLM causal-judgement and narrative-retention protocol

We implemented a model-side protocol corresponding to the human behavioral experiment using textbased descriptions of the same 50 predefined events. This adaptation preserved the event structure and causal logic of the narrative while presenting the stimulus in the model’s native modality. Although the resulting text inputs were not perceptually identical to the audiovisual stimulus viewed by human participants, using text reduced the influence of low-level audiovisual features and allowed us to focus on the causal organization of the narrative.

To reproduce the online constraint of the human task, causal-role judgements were generated incrementally. For event *i*, the model received only the descriptions of events 1 through *i*, followed by the fixed instruction:

*Given the story so far (events 1 through i), what is the primary function of event i? Please choose one: (i) no clear bridging function; (ii) explains previous events; (iii) sets up subsequent developments; or (iv) uncertain*.

This incremental presentation prevented the model from accessing narrative events occurring after event *i*, thereby matching the information available to participants during online causal judgement. After processing the complete narrative, the model received the following instruction:

*Write a short and coherent summary of the story you have just read, including as many key plot points as possible, without listing event numbers*.

To reduce potential contamination from film-related content encountered during pretraining, all person and place names were replaced with generic identifiers such as personA and placeA in the LLM event descriptions. This replacement was applied only to the LLM inputs; human participants viewed the original, unmodified audiovisual stimulus. Further analyses of this LLM-specific control are reported in Supplementary Information 2.

#### Cross-system event-wise retention measures

Human recall summaries were transcribed and segmented into clauses or short utterances, whereas LLM-generated summaries were segmented into corresponding textual units. Both outputs were aligned to the same 50 predefined events to determine whether each event was mentioned. Human recall richness was defined as the proportion of participants who mentioned a given event in their recall summary, whereas model-side retention was represented by the corresponding event-wise mention measure.

Both tasks involved narrative summarization, with human summaries produced from memory after viewing and LLM summaries generated with the complete narrative available in context. Narrative summaries tend to preserve gist, macrostructure and structurally central information [43, 44]. Aligning the two outputs to the event units used in the Sherlock fMRI analysis therefore provided an approximate shared event-level output space for comparing the narrative information included in human recall summaries and LLM-generated summaries.

### 4.3 Reactivation algorithm

To quantify *neural reactivation* during naturalistic narrative processing, we implemented a boundary-triggered representational-similarity analysis following the approach introduced by Hahamy et al. [7].

To account for the canonical haemodynamic delay, boundary time points were shifted by 4 TRs (6 s; TR=1.5 s; Sherlock dataset already includes 3TRs shift and adds another TR) prior to pattern extraction.

#### Event-boundary and event representations

For each participant, we extracted multivoxel patterns within a whole-brain searchlight (sphere radius: 3 voxels, 9 mm). Event-boundary representations were defined as the voxel pattern in the single TR marking the end of each event. Event representations were defined as the mean voxel pattern across within-event TRs that were at least 5 TRs (7.5 s) away from adjacent boundaries, to minimize boundary-related contamination.

#### Boundary–event similarity matrix

Within each searchlight, we computed the Pearson correlation between each event-boundary representation and each event representation, yielding an event-boundary × event similarity matrix *S*∈ ℝ^*N ×N*^, where *N* = 42 after excluding short events from the original 50-event segmentation. Correlation coefficients were Fisher *z*-transformed before summary. The main diagonal of **S** (boundary *i* vs. the immediately preceding event *i*) was removed to focus on temporally remote events.

#### Reactivation index

Because between-event similarity can produce symmetric correlations with both past and future events, we computed a *reactivation index* that contrasts boundary similarity to past versus future events, cancelling out time-symmetric scene-similarity effects. Let *P* = {(*i, j*) : *j < i*}denote the past (lower-triangular) entries and *F* = {(*i, j*) : *j > i* }denote the future (upper-triangular) entries (excluding removed diagonals). The reactivation index was defined as

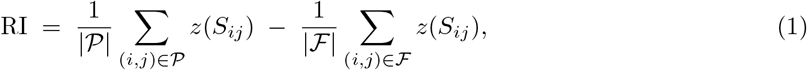

where *z*(·) is the Fisher transform. Following prior work, we additionally discarded a matched set of near-diagonals from both the past and future halves of **S** to reduce temporal-autocorrelation artefacts induced by preprocessing filters.

#### Reactivation-strength matrix

In addition to the primary signed reactivation index, we constructed an unsigned *reactivation-strength matrix* for descriptive visualization of the overall magnitude of boundary–event representational coupling.Starting from the event-boundary × event similarity matrix **S**, we applied the Fisher *z*-transform to each Pearson correlation coefficient and then took its absolute value, yielding a matrix **R** ∈ ℝ^*N ×N*^ with entries

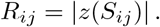

Each element of **R** therefore indexes the magnitude of coupling between the representation at the boundary of event *i* and the representation elicited during event *j*, irrespective of the sign of the original correlation. Importantly, this unsigned matrix was used only to visualize and compare the overall distribution of boundary–event coupling across events, participants and model layers; it was not used as the primary inferential measure of reactivation. Because negative correlations do not by themselves provide unambiguous evidence for reinstatement, all statistical tests of causal reactivation were performed on signed Fisher-*z*-transformed similarities or on the signed past-minus-future reactivation index. We further verified that the causal reactivation effects were robust when using signed similarity values, after excluding the main diagonal and matched near-diagonal entries to reduce temporal-autocorrelation artefacts, and after controlling for time-symmetric background scene similarity arising from shared semantic and contextual content, through the past–future similarity contrast and for local scene activity through pre- and post-boundary comparisons (Supplementary Information 9). Thus, the unsigned reactivation-strength matrix should be interpreted as a descriptive measure of coupling magnitude, whereas the signed reactivation index provides the basis for statistical inference about temporally asymmetric, boundary-triggered reactivation.

### 4.4 Causal-role-specific reactivation analysis

#### Construction of causal-role masks

behavioral causal judgements were aggregated across participants using the Dawid–Skene model described in Section 4.1. The resulting event-level estimates were used as behavioral reference annotations. Each event was assigned to one of four causal-role categories: *Result, Cause, Independent* or *Other*.

These assignments were used to construct the four masks shown in Fig. 3b, with each boundary labelled according to the event that had just ended.

For each causal-role category *c*, its reactivation index was obtained by applying the corresponding mask *M* ^(*c*)^ to the directional reactivation matrix. In general form, the category-specific index was calculated as

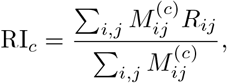

where *R*_*ij*_ denotes the directional reactivation value relating event *i* to the boundary of event *j*. The precise calculation of *R*_*ij*_, including the subtraction of the reverse-direction similarity, is described in Section 4.3. Indices were calculated separately for each participant in the fMRI data and for each layer in the LLM.

#### Voxel-wise causal contrasts

To identify brain regions in which reactivation differed across causal roles, category-specific reactivation indices were calculated independently at each voxel or searchlight centre. For any two causal categories *A* and *B*, the contrast map was defined as

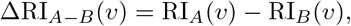

where *v* denotes the voxel or searchlight centre. Positive values indicate greater reactivation for category *A*.

Group-level inference for the Result–Cause contrast was performed using two-sided paired (t)-tests, with statistical significance assessed at (P¡0.05) after family-wise error (FWE) correction. The Result–Cause contrast was designated as the primary directional contrast because it directly tested whether a causal outcome boundary preferentially reinstated representations associated with preceding causal events. The remaining contrasts assessed whether the observed effect generalized to, or could be distinguished from, differences involving Independent and Other events.

#### Parcel-level localization

To characterize regional differences in reactivation coverage, we compared the corrected Result–Cause contrast map with our replication of the original reactivation analysis, which did not distinguish events by causal role. Both thresholded maps were summarized using the Schaefer 400-parcel cortical atlas [27]. For each parcel *k*, we calculated the proportion of significant voxels in each map:

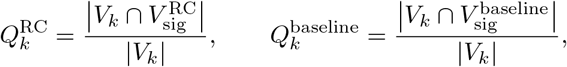

where *V*_*k*_ denotes the set of voxels in parcel *k*, and 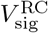 and 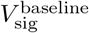 denote the sets of voxels meeting the significance criteria for the Result–Cause contrast and the replicated original reactivation analysis, respectively. The regional difference in significant-voxel proportions was defined as

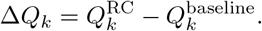

Parcels were ranked by Δ*Q*_*k*_, and the ten parcels with the largest positive differences were displayed in Fig. 3i. Normalizing by parcel size allowed comparison of significant-voxel proportions across parcels of different sizes. This analysis descriptively compared regional coverage between the two thresholded maps; no additional inferential test of the between-map differences was performed.

The other causal-role contrasts served to evaluate the specificity of the primary Result–Cause effect and are reported in the voxel-wise analysis.

#### Meta-analytic spatial comparison

NeuroSynth is a platform for automated meta-analysis of published fMRI studies that provides brain maps associated with terms in the neuroimaging literature [28]. We used the whole-brain Result–Cause contrast map as input and calculated its spatial Pearson correlations with NeuroSynth term-based maps to explore the functional associations of the observed reactivation pattern.

### 4.5 Estimation and evaluation of event-to-boundary reconstruction coefficients

We estimated an event-to-boundary reconstruction matrix separately for each human participant and each LLM layer. The model quantified how preceding event representations jointly accounted for each boundary representation. The fitted coefficients were then evaluated against the event-pair causal annotations.

Concretely, let *E* ∈ ℝ^*n×d*^ contain the retained event representations and *B* ∈ ℝ^*n×d*^ the corresponding boundary representations, with events ordered chronologically. Each row *e*_*i*_ or *b*_*i*_ is a *d*-dimensional representation. Event and boundary extraction followed the dataset-specific definitions described above. We modelled the boundary representations as

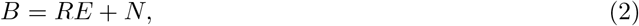

where *R*_*ij*_ is the coefficient of event *j* in the reconstruction of boundary *i*, and *N* contains the residuals.

The matrix *R* summarizes the linear relationships between event and boundary representations. Under these assumptions, estimating replay reduces to the constrained least-squares problem

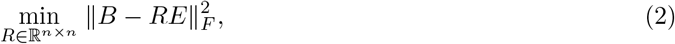

where∥·∥ _*F*_ denotes the Frobenius norm. Because each event boundary can depend only on events that have already occurred, *R* is naturally lower-triangular, which allows us to solve for it row by row.

Writing *E* = [*e*_1_, …, *e*_*n*_]^*⊤*^ and *B* = [*b*_1_, …, *b*_*n*_]^*⊤*^, we estimate the coefficients *R*_*i*,1:*i™*1_ by regressing the boundary vector *b*_*i*_ on the preceding event vectors *e*_1_, …, *e*_*i™*1_,

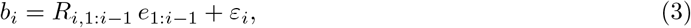

which yields the closed-form least-squares solution

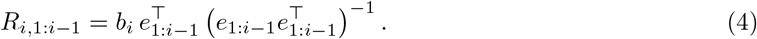

Here *R*_*i*,1:*i™*1_ denotes the coefficients in row *i* and columns 1 through *i* ™ 1 of the reconstruction matrix, and *ε*_*i*_ is the residual for event boundary *i*. Intuitively, the replay coefficients *R*_*i*,1:*i™*1_ describe how strongly each past event contributes to the boundary at time *i*, and together the matrix *R* serves as a replay matrix that we can compute for both human data and LLM activations, then compare as an abstract signature of causal-structure learning.

Now we will show how to calculate the replay matrix. We denote reply matrix as R, react matrix as A, B and E, as above, represent event-boundary matrix, and event matrix, respectively. From the calculations of react matrix and reply matrix above, we can see that:

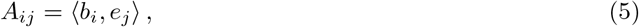

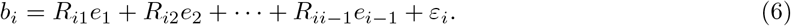

We perform an inner product of both sides of the expression 6 with respect to *e*_*j*_, *j* ∈ [1, *n*], and the event similarity matrix be S, where *S*_*ij*_ = ⟨*e*_*i*_, *e*_*j*_⟩, then we can get:

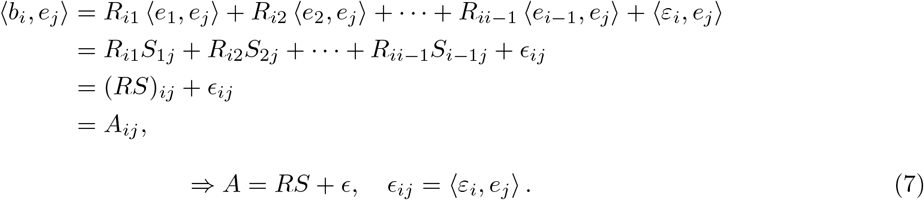

Let’s assume the event similarity matrix is invertible, so the above formula means :

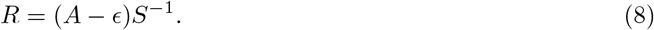

This indicates that the replay matrix *R* is derived by filtering extraneous noise from the reactivation matrix *A* and subsequently computing a weighted aggregation using the event similarity matrix *S*. Moreover, it indicates that the replay matrix captures causal learning dynamics at a finer granularity, enabling more detailed inference of the intelligent system’s underlying causal acquisition process.

### 4.6 Voxel-wise probe

We used a voxel-wise deletion analysis to identify brain voxels contributing to the similarity between brain and LLM event-to-boundary reconstruction matrices.

For participant *s*, events with entirely missing event or boundary representations were excluded. Voxels containing missing values across the retained events were also excluded. The same event mask was applied to the LLM representations, ensuring that brain and LLM reconstruction matrices used identical event coordinates.

Using the procedure in Section 4.5, we first computed the brain reconstruction matrix from all valid voxels. For LLM layer*l*, the baseline discrepancy was

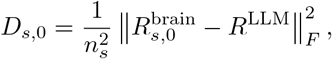

where *n*_*s*_ is the number of retained events.

We then removed voxel *v* from both the event and boundary representations, repeated the row-wise normalization, and recomputed the brain reconstruction matrix. The discrepancy after deletion was

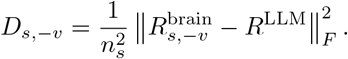

Each deletion started from the full set of valid voxels. The reconstruction mask and LLM reference were held fixed.

The contribution of voxel *v* was defined as

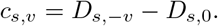

Positive values indicate that removing the voxel weakened brain–LLM alignment. Larger positive values therefore identify voxels whose removal disrupted the correspondence more strongly.

For each participant and layer, we ranked positive contribution scores in descending order and selected up to 20% of the valid voxels. Specifically, the selection budget was

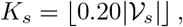

where *V*_*s*_ is the participant’s valid voxel set. Continuous contribution scores and selection masks were mapped back to the original voxel coordinates.

### 4.7 Preprocessing and Manifold Statistics Extraction

For each subject and condition, we represent the neural trajectory as a sequence of high-dimensional response vectors. All trajectory points are standardized using z-score normalization. The standardized high-dimensional responses are used for the subsequent manifold-statistics analysis.

For visualization, the standardized responses are projected to a six-dimensional principal component space using PCA and further embedded into three dimensions using UMAP with cosine distance, spectral initialization, and densMAP regularization. The UMAP model is fitted on the sampled trajectories and then applied to all trajectories using the learned transformation. After embedding, the concatenated low-dimensional points are split back into individual trajectories according to their original temporal lengths, yielding one three-dimensional point cloud for each trajectory. Thus, dimensionality reduction is used for visualization rather than for the manifold-statistics analysis.

To quantify the topological structure of each trajectory manifold, we construct an alpha complex from the standardized high-dimensional representations and compute persistent homology from the resulting simplex tree. Persistence intervals are extracted for both connected components (*H*_0_) and one-dimensional cycles (*H*_1_). Each persistence diagram is converted into birth-persistence coordinates by replacing death time with persistence length. Infinite intervals are removed before vectorization.

Finally, the *H*_0_ and *H*_1_ diagrams are separately transformed into persistence images with resolution (15× 15) and Gaussian bandwidth (0.5). The two persistence images are flattened and concatenated to obtain a fixed-length topological feature vector for each trajectory. These vectors provide a compact statistical representation of the manifold topology and are used for subsequent comparison across subjects, conditions, and train-test splits.

## Acknowledgements

The research in this article is supported by the New Generation Artificial Intelligence of China (2024YFE0203700), National Natural Science Foundation of China under Grants U22B2059 and 62576124.

